# Division Asymmetry Drives Cell Size Variability in Budding Yeast

**DOI:** 10.1101/2025.10.22.683920

**Authors:** Félix Proulx-Giraldeau, Xin Gao, Yagya Chadha, Jordan Xiao, Kurt M. Schmoller, Jan M. Skotheim, Paul Francois

## Abstract

Cell size variability within proliferating populations reflects the interdependent regulation of cell growth and division as well as intrinsically stochastic effects. In budding yeast, the G1/S transition exerts strong size control in daughter cells, which manifests as the inverse correlation between how big a cell is when it is born and how much it grows in G1. However, mutations affecting this size control checkpoint only modestly influence population-wide size variability, often altering the coefficient of variation (CV) only by ∼10%. To resolve this paradox, we combine computational modeling and live-cell imaging to identify the principal determinants of cell size variability. Using an experimentally validated stochastic model of the yeast cell cycle, we perform parameter sensitivity analysis and find that division asymmetry between mothers and daughters is the dominant driver of CV, outweighing the effects of G1/S size control. Experimental measurements across genetic perturbations and growth conditions confirm a strong correlation between mother-daughter size asymmetry and population CV. These findings reconcile previous observations and show how asymmetric division operates in concert with G1/S size control to govern cell size heterogeneity.

## Introduction

Cell size is intrinsically linked to cellular function. For example, red blood cells must be small enough to efficiently move through capillaries, while macrophages need to be large enough to engulf pathogens. As cells grow larger, the concentration of biochemical components changes to alter the internal biochemistry of cells. If cells become excessively large, they grow poorly and exhibit features of senescence (Crozier et al., 2023; Foy et al., 2023; Khurana et al., 2023; Lanz et al., 2022, 2024; Manohar et al., 2023; Miettinen & Björklund, 2016; Tzur et al., 2009; Zatulovskiy et al., 2022). To avoid such a detrimental outcome, organisms have evolved robust regulatory circuits that sense and control cell volume across successive divisions, ensuring that biomolecular concentrations remain within functional limits. These size-control mechanisms have been extensively reviewed, providing insight into how cells link their growth and division cycles to preserve homeostasis in diverse contexts (Ginzberg et al., 2015; Lloyd, 2013; Xie et al., 2022).

One way eukaryotic cells actively control their size is by linking cell growth to specific cell cycle transitions. For example, in fission yeast a sharp cell size threshold governs entry into mitosis (Fantes, 1977; Sveiczer et al., 1996). In contrast, budding yeast shows its strongest size control at the G1/S transition in its smaller daughter cells (Di Talia et al., 2007, 2009; Johnston et al., 1977). The strength of a size control mechanism can be quantified by plotting the amount of cell growth during a given phase against the size the cell was when it entered that phase. A strong negative correlation (slope near –1) indicates that initial size differences are nearly fully corrected within one division cycle as is the case for both fission yeast and mouse epidermal stem cells (Fantes, 1977; Sveiczer et al., 1996; Xie et al., 2024; Xie & Skotheim, 2020). The corresponding slope for budding yeast at the G1/S transition is lower but still appreciable (Di Talia et al., 2007). Many bacteria employ a weaker form of size control called an “adder,” in which each cell adds a nearly constant volume from birth until division (slope near 0), regardless of its initial size (Amir, 2014; Eun et al., 2018; Jun et al., 2018; Westfall & Levin, 2017; Willis & Huang, 2017). In symmetrically dividing species, this adder mechanism gradually pushes cells towards a target size over multiple generations.

While measuring the correlations between size and growth can be used to assess size control in specific cell cycle phases, a more holistic way to assess cell size control is to consider the entire distribution of cell sizes. In this view, mutations affecting size control are defined as those that alter the coefficient of variation (CV), which is the ratio of the standard deviation to the mean (CV = *σ*/*μ*). Because this metric is normalized, it reveals how tightly cell sizes cluster irrespective of the average cell size, making it possible to compare cells across different conditions or even different organisms. Surprisingly, recent work demonstrated that budding yeast mutations with large effects on cell size only modestly or even imperceptibly affect the CV (Barber et al., 2020; Chen et al., 2020). In other words, while the average cell size changes in response to these mutations, the shape of its distribution does not. This observation raises the question of whether we have fundamentally misunderstood cell size control. Specifically, what connection should we anticipate between mutations and population-wide size variability, and is our current understanding of how growth is linked to specific cell cycle transitions compatible with these unexpected findings?

Here, we use both experimental and theoretical approaches to investigate how cell size control mutations influence population size variability. We employ an established computational model for the budding yeast cell cycle featuring a size-dependent G1/S transition in daughter cells, followed by more weakly regulated S/G2/M phases (Chandler-Brown et al., 2017). Through parameter sensitivity analysis, we find that the primary driver of changes in the CV is the division mechanism itself. Namely, the CV is primarily controlled by the difference in cell sizes at division between the larger mother and smaller daughter cells. Altering mother–daughter asymmetry proportionally shifts the CV as predicted by our computational model. In addition, G1/S size control mutations typically modulate the CV by 10–20%, consistent with the measured effects of size control mutations affecting G1/S control. Our findings show how the current framework for yeast growth and division can reconcile the paradox of why size control mutations have a limited impact on CV.

## Results

Recent work showed that mutations affecting budding yeast cell size only modestly or even imperceptibly affect the CV of the population (Chen et al., 2020). A typical change in the CV was about 10% even when average cell size increased as much as 50% (**Fig. 1**). This raised the question as to how sensitive we should expect the population CV to be to mutations that affect cell size? To determine how sensitive the CV is to mutations to parameter values, we turned to a mathematical model of the growth and division of budding yeast cells that was previously validated experimentally (Chandler-Brown et al., 2017). While various molecular models have been proposed to explain the size-dependence of the budding yeast cell cycle (Barber et al., 2020; Schmoller et al., 2015), the Chandler-Brown model is agnostic to this debate and coarse-grains the molecular machinery into the physiological dependencies of the different cell cycle transitions. In this model, each cell cycle transition is determined by a stochastic rate that depends on physiological parameters such as cell size or growth rate. While this model is simple enough to comprehensively analyze, it is complex enough to accurately describe progression through the budding yeast cell cycle in both the larger mother and smaller daughter cells following cell division (**Fig. 2A**). The model is described in the **Materials and Methods** section. This model has 21 independent parameters that were originally fit to WT data and strikes a good balance between phenomenology and descriptiveness (Chandler-Brown et al., 2017). Notably, G1 phase is broken up into a pre-*Start* and a post-*Start* phase, where the *Start* transition marks the point of commitment to the cell cycle with respect to exposure to mating pheromone, which coincides with the export of the cell cycle inhibitor Whi5 from the nucleus (Hartwell et al 1974; Skotheim et al 2008; Charvin Siggia Cross 2010; Doncic Skotheim 2011). Cell size control is implemented at *Start* in daughter cells mainly via a linear size-dependent rate of cell cycle progression. In this way, cell size control in this model is stochastic and thus imperfect. In addition, mother cells accumulate size over successive generations because they have a non-zero growth phase in G1 and cannot lose this accumulated mass because the daughter cell is only made from the mass accumulated in the S/G2/M phases of the cell cycle. Because growth is close to exponential during most of the cell cycle, a bigger cell will accumulate mass faster than a smaller cell. This has an important consequence which will become relevant later on, namely, that a bud grown on an older cell (2nd generation or older) will become a larger daughter cell than if its progenitor was a 1st generation cell (**Fig. 2A**).

**Figure 1:**
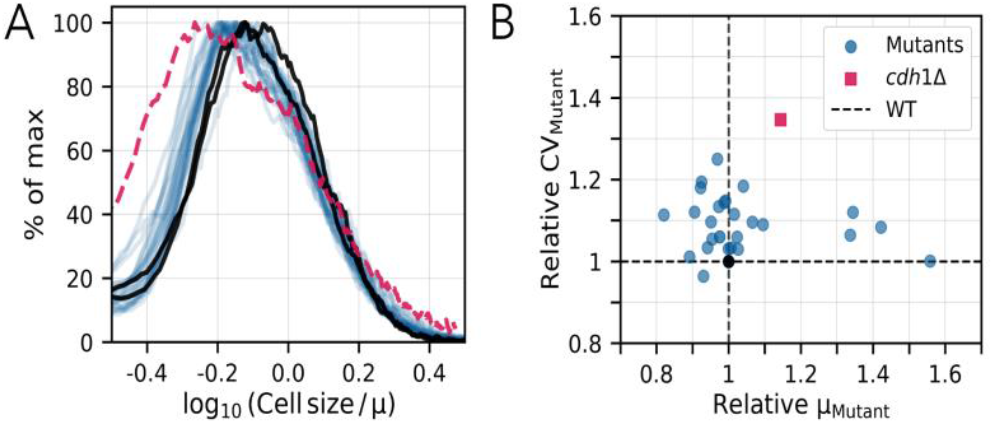
Mutations affecting cell size have a modest effect on the coefficient of variation of size within the population. **(A)** Distributions of cell size after normalizing by the population mean cell size for 2 WT (black) and 29 mutant strains (colored) of the budding yeast *S. cerevisiae*. Data are from (Chen et al., 2020). **(B)** The coefficient of variation (CV_Mutant_ = standard deviation/mean) for each size mutant normalized by the CV of WT cells plotted against the average cell size for each mutant (µ_Mutant_) population normalized to the average size of WT cells. Red dashed curve in panel A and red square in panel B corresponds to the *cdh1Δ* mutant with the largest change in CV compared to WT.

**Figure 2:**
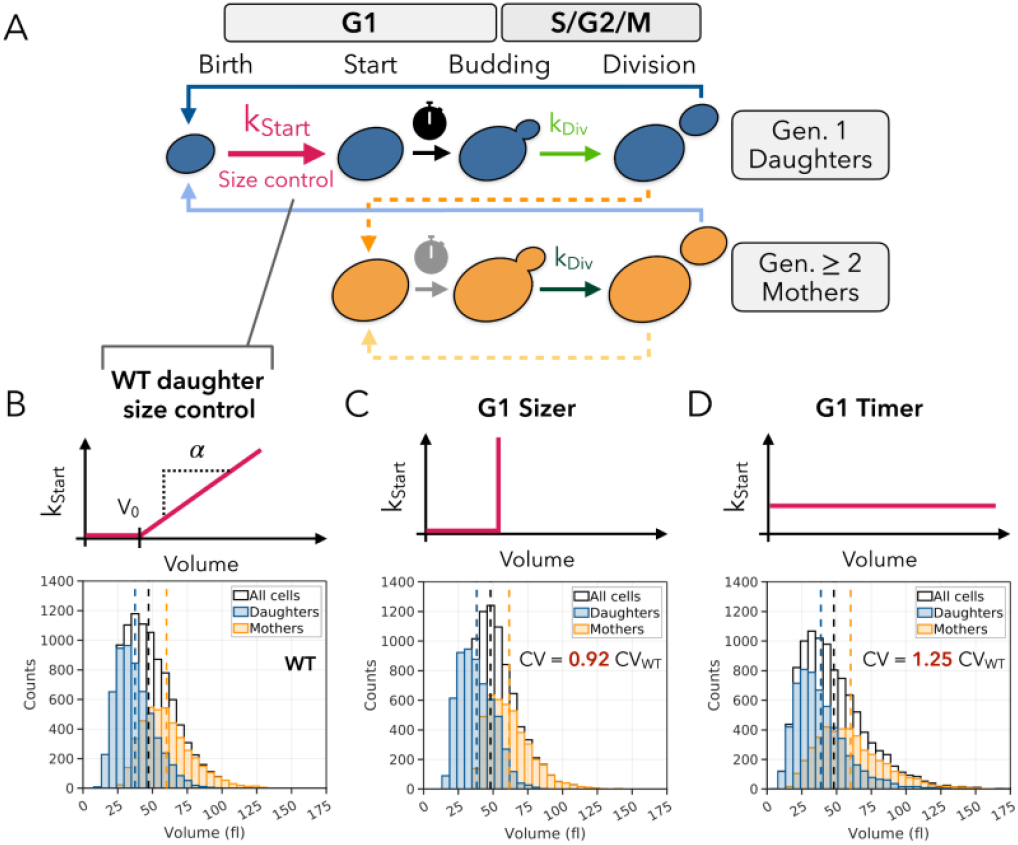
A mathematical model of the budding yeast cell cycle indicates that a ∼10% change in CV in response to a cell size mutation is typical. **(A)** Schematic diagram of the model constructed by (Chandler-Brown et al., 2017) to describe progression through the asymmetric budding yeast cell cycle. **(B)** WT version of *Start* size control where the stochastic rate of progression through the transition depends linearly on the current cell size above a threshold V_0_. The resulting steady-state cell size distribution with model parameters fit to WT data is shown below. For comparison with simulations below, we find CV_WT_ = 0.41 **(C)** The *Start* transition is modified to be a deterministic sizer where cells instantaneously progress in the cycle once they reach a threshold size. The resulting steady-state size distribution is shown below with a CV = 0.92 CV_WT_. **(D)** The *Start* transition is modified to take place at a constant stochastic rate, resulting in an effective pre-*Start* timer with some noise due to stochasticity. The resulting steady-state size distribution is shown below with a CV = 1.25 CV_WT_. In (C)-(D) all other aspects of the cell cycle model are kept constant.

### The population CV responds moderately to the drastic perturbations of *Start*

To begin our analysis of the sensitivity of the CV of cell size distributions, we first set out to perturb the cell size control mechanism at *Start* and then see how this affected the CV of the population cell size distribution at steady state. We chose this metric of variability because it is the most basic measurement of a complex population of mother and daughter cells at all phases of the cell cycle. Moreover, the distribution and population CV are easily accessible experimentally using either a Coulter counter or flow cytometer (Berenson et al., 2019), which is why it is commonly used in publications analyzing the effects of cell size control (Barber et al., 2020; Chen et al., 2020).

The implementation of the size control mechanism in the Chandler-Brown model is an imperfect sizer where the rate of going through the *Start* checkpoint increases linearly with cell size at a slope *α* and offset *V*_0_ (**Fig. 2B**). One drastic change of this mechanism could be to turn it into a perfect sizer mechanism, in which the *Start* checkpoint becomes a deterministic transition once cell size reaches a threshold size. This can be implemented easily in the model by setting the slope *α* to ∞. We then fixed the free parameter *V*_0_ such that the average size of the resulting steady-state distribution would be identical to that of WT, thus avoiding any potential change to CV solely due to a change in average size. With a perfect sizer mechanism operating at the *Start* transition, we measure a relative change in CV of -8% (**Fig. 2C**). Conversely, we can abolish the size-sensing mechanism by setting the slope *α* to 0. Specifically, by setting the rate of progression through *Start* to a constant rate independent of size, we can turn the pre-*Start* G1 into a stochastic timer where cells spend on average the same time in pre-*Start* G1 irrespective of their initial size at birth. If we set this constant rate of progression through *Start* such that the average size of the resulting steady-state distribution remains unchanged from WT, we find that the CV increases by 25% (**Fig. 2D**).

So far, our analysis suggests that even drastic perturbations to cell size control at *Start* in *S. cerevisiae*, where parameters are either set to 0 or ∞, only yield relative changes in CV up to 25%. In this light, the changes to the CV of the mutants shown in **Fig. 1** lie within an expected small range (0-20%) of what one might expect from a significant perturbation to size control at *Start*. In other words, we should not expect to see any 50% to 100% changes to the CV from any perturbations affecting the cell cycle in G1.

### Live cell analysis of *Start* mutants is consistent with moderate changes to the CV

Next, we sought to experimentally test our new intuition for how the population CV should respond to mutations of genes known to regulate *Start*. Namely, we predict that the effect on CV should be modest and below 20%. We chose to analyze two of the most well-known mutants of the *Start* checkpoint, *cln3Δ* and *cln3Δwhi5Δ* (Bruin et al., 2004; Costanzo et al., 2004; Cross, 1988; Nash et al., 1988). Cln3 is an activator while Whi5 is an inhibitor of *Start* and both proteins have been linked to specific size control mechanisms (Qu et al., 2019; Schmoller et al., 2015, 2022; Wang et al., 2009).

To identify the effect of mutations to known *Start* regulators on the population CV, we imaged unperturbed cells growing and dividing asynchronously in the same conditions as used by (Chandler-Brown et al., 2017). The cells were grown in a synthetic complete medium using glycerol and ethanol as the carbon source. In these conditions, cells have a longer G1 phase and exhibit stronger size control at *Start* in daughter cells compared to growth in glucose medium (Di Talia et al., 2007). Cells were then segmented and tracked using the Cell-ACDC software (see Methods) (Padovani et al., 2022). This allowed us to measure the cell size of 377 *cln3Δ* cells and 504 *cln3Δwhi5Δ* cells at steady state (**Fig. 3C, E**). The CV increased ∼9% for *cln3Δ* mutant cells and ∼27% for *cln3Δwhi5Δ* double mutant cells. It is notable that removing the size-sensing inhibitor Whi5 along with the activator Cln3 yields cells with an average size similar to WT cells, yet with a CV of 27% bigger than the WT. This is approximately the magnitude of change we expected from turning the WT size control mechanism into a stochastic timer (**Fig. 2D**).

**Figure 3:**
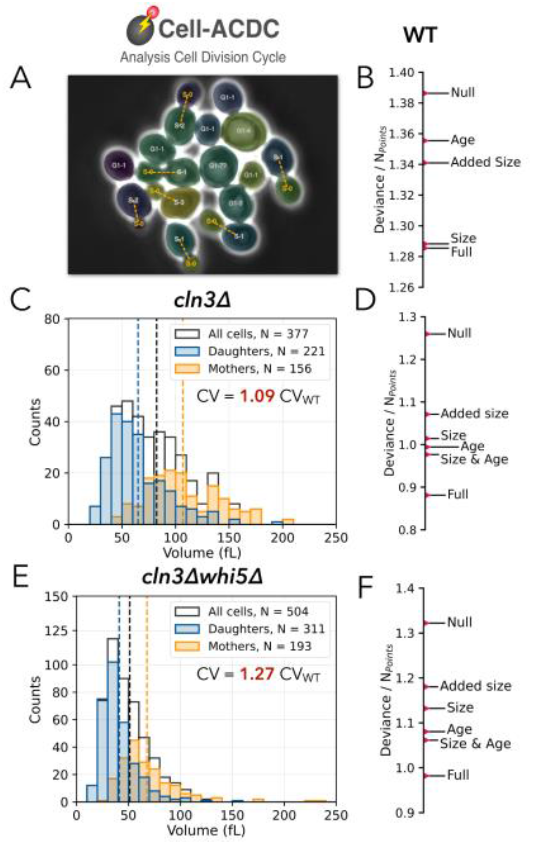
Logistic regression models predicting passage through *Start* for mutant cells lacking key G1 regulators. **(A)** Cell-ACDC (Padovani et al., 2022) was used to segment, track and analyze the cell cycles of the live cell mutants shown in panels. Phase contrast is used to identify cell boundaries. The budneck is marked with the Myo1-mKate fluorescent protein to detect cell division. **(B)** Deviance measurements for logistic regressions of progression through *Start* based on the indicated predictors show that cell size is the most informative single predictor for this transition. Data from (Chandler-Brown et al., 2017). In (B),(D) and (F), *Full* indicates logistic regression using all 3 predictors. *Null* indicates the control model with no predictors. **(C)** Steady-state cell size distribution of *cln3Δ* daughter and mother cells at all phases of their cell cycle, n = 377, CV = 1.09 CV_WT_. **(D)** Deviance measurements for indicated logistic regressions predicting the passage through *Start* of *cln3Δ* mutant cells. **(E)** Steady-state size distribution of *cln3Δwhi5Δ* daughter and mother cells at all phases of their cell cycle, n = 504, CV = 1.27 CV_WT_. **(F)** Deviance measurements for indicated logistic regressions of predicting passage through *Start* of *cln3Δwhi5Δ* mutant cells.

Next, we set out to investigate whether size was still the best predictor of the *Start* transition in the *cln3Δ* and *cln3Δwhi5Δ* mutants as had previously been found for WT cells (Chandler-Brown et al., 2017). To test this, we performed a series of logistic regressions using individual and combinations of statistical predictors to describe the *Start* transition. Following Chandler Brown et al. (2017), we chose cell size, age, and added size since birth as the individual predictors of the timing of the transition. We also combined predictors together to perform 2- and 3-variable logistic regressions. We then examined the deviance of the prediction for each logistic regression to identify the best predictors of the transition. The deviance can be interpreted as a measure of goodness of fit: the lower the deviance, the more accurate the logistic regression is at predicting the *Start* transition. The resulting deviance measurements for the WT data of the original publication are shown in **Fig. 3B** with cell size being the best individual predictor of *Start* in this case. In both the *cln3Δ* (**Fig. 3C-D**) and *cln3Δwhi5Δ* (**Fig. 3E-F**) mutants, we see cell size become less important as an individual predictor in favor of cell age. This is particularly interesting given the expected magnitude of change of CV for a pure stochastic timer in pre-*Start* G1 and the resulting change in CV for this mutant (**Fig. 2D**). Indeed, a stochastic timer is a good description of the pre-*Start* G1 phase of *cln3Δwhi5Δ* mutant daughter cells.

### Parameter sensitivity analysis identifies division mother-daughter asymmetry as a major determinant of cell size variability

After finding that mutations affecting the cell size control mechanism at the *Start* checkpoint had only a limited impact on the CV of the steady-state size distribution, we next sought to identify which (other) cell cycle parameters exert a more significant influence. To do this, we went back to theory and performed a local parameter sensitivity analysis of the Chandler-Brown model by computing the numerical gradient of the CV in the 21-dimensional parameter space of the model (**Fig. 4A**, see details in **Materials and Methods**). This gradient is a 21-dimensional vector that quantifies the local contribution of each parameter to the CV, enabling direct comparisons between them. Because parameters have different units and scales, we need to rescale them properly for comparison. To do this, we followed a standard strategy and computed the numerical gradient with respect to the logarithm of each parameter which can be interpreted as transforming every parameter into a free-energy equivalent (Gutenkunst et al., 2007). Additionally, we examined both the CV of the steady-state distribution (CV_SS_) and that of the daughter size distribution at the *Start* checkpoint (CV_*Start*_). While CV_*Start*_ is less readily accessible experimentally because it requires live-cell tracking, it is particularly relevant to the previously studied cell size control mechanisms operating at the *Start* transition in daughter cells. Since the resulting 21-dimensional gradient vectors for both CVs are difficult to interpret at first glance, we sought to gain an intuition for how these perturbations affect cell size control by perturbing parameters along the gradient direction and observing the resulting effects on cell cycle dynamics. Specifically, we explored the gradient’s influence by modifying the value of each individual parameter in the direction of the gradient. The i^th^ parameter *θ*_*i*_ becomes 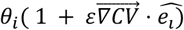, where *ê*_*i*_ is a unit vector whose sole component is in the direction of parameter *θ*_*i*_, and where *ε* = 0.5 is a scaling factor. Since the gradient identifies the direction in parameter space where the magnitude of the CV changes the most, we define positive *ε* as the direction yielding an increase in CV.

**Figure 4:**
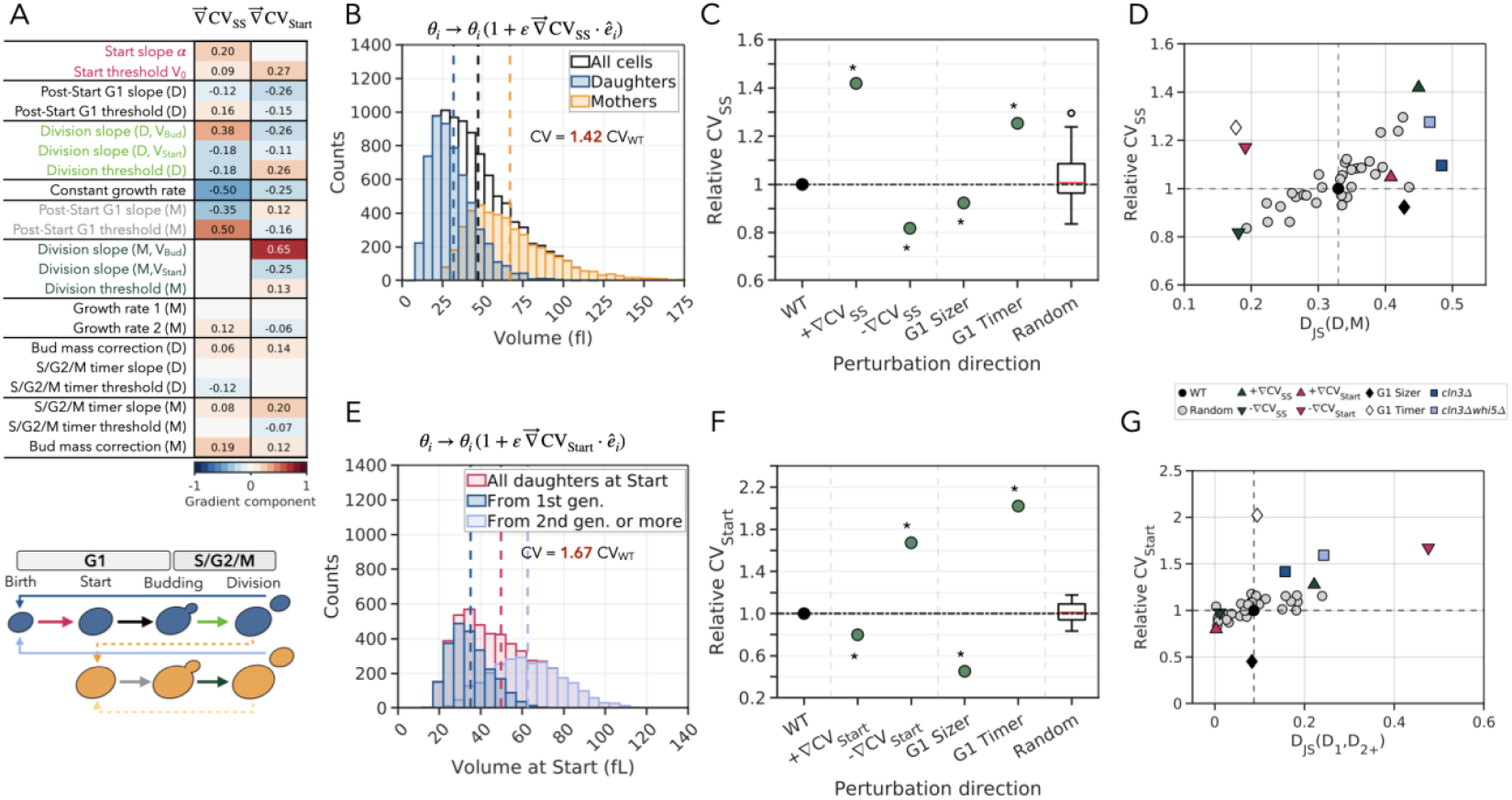
Parameter sensitivity analysis of the model identifies two types of asymmetry as the main drivers of the CV of the steady-state cell size distribution and of the daughter cell size distribution at the *Start* transition. **(A)** Vectors representing the numerical gradients with respect to the logarithm of the parameters of the model. The first column represents the gradient of the CV of the steady state size distribution, and the second column represents the gradient of the CV of the daughter size distribution at the *Start* transition. Rows indicate each parameter’s contribution to the gradients as indicated by their names on the left, with colors representing the magnitude of each parameter’s contribution to the gradient. Below, a schematic representation of the cell cycle model is shown where colored arrows indicate the stochastic rates of transition with colors matching the names of the parameters on the rows above. **(B)** Resulting steady-state size distribution of mother and daughter cells upon perturbing the model parameters along the direction of the gradient of the CV of the same distribution. *ε* = 0.5, CV = 1.42*CV_WT_. **(C)** CV of the steady-state size distribution relative to the WT value indicated by the first black dot. The following 4 green dots correspond to parameter perturbations of the model, first along the positive and negative directions of the gradient as is done in panel (B) with *ε* = ±0.5, then by changing the G1 size control mechanism to a pure sizer or timer as is done in Fig. 2C,D. The boxplot represents the quartile distribution of relative CV when randomly and uniformly perturbing parameters with the same magnitude as before with *ε* = 0.5. Stars indicate statistical significance p<0.01 after performing a paired sample t-test between the directed parameter perturbations and the random parameter perturbations. **(D)** Relative CV of the steady-state size distribution of cells plotted against the asymmetry of the mother and daughter sub-distributions as quantified using the Jensen-Shannon divergence. Data of previous panels of this figure along with the experimental data for the *Δcln3* and *Δcln3Δwhi5* mutant cells shown in Fig. 3C,E. **(E)** Resulting daughter size distribution at the *Start* transition upon perturbing the parameters along the direction of the gradient of the CV of the same distribution. *ε* = 0.5, CV = 1.67 CV_WT_. **(F)** CV of the daughter size distribution at the *Start* transition relative to the WT value, similar to panel (C). **(G)** Relative CV of the daughter size distribution at *Start* against the asymmetry of the daughters born from 1^st^ generation cells and the daughters born from 2^nd^ generation or later cells sub-distributions as indicated by the Jensen-Shannon divergence. Data of previous panels of this figure along with the experimental data for *Δcln3* and *Δcln3Δwhi5* mutant cells shown in Fig. 3C,E.

From our computation (**Fig. 4A**), we see that the dominating directions in parameter space driving changes to the steady state size distribution, identified by the ^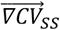^, can be explained to be the combination of two main effects: decreasing the growth rate and lengthening the post-*Start* G1 timer for the mothers. Slow growth decreases the size of the bud, resulting in a decreased size at birth for new daughter cells. While slowing down the growth rate creates smaller cells in general, this effect is less in mother cells because they also lengthen their post-*Start* G1 phase. Altogether, this parameter direction corresponds to a regime where mothers are bigger and daughters are smaller, which is thus expected to amplify the asymmetry between these two types of cells to widen the overall cell size distribution with an increase of CV of more than 40% in our simulations for *ε* = 0.5 (**Fig. 4B)**. Our gradient analysis further confirms the modest effect of *Start* size control parameters on CV relative to the first two vector components shown in **Figure 4A**. Importantly, modifying parameters along the gradient direction has a greater effect on CV than altering the G1 size control mechanism to a pure sizer or timer, or randomly perturbing the parameter space with the same magnitude (**Fig. 4C**). This is noteworthy because shifting the control mechanism requires extreme parameter modifications, such as setting a parameter to zero or infinity, whereas perturbations along the gradient direction are comparatively subtle, yet more effective in modulating the CV.

Having identified daughter-mother asymmetry as crucially important for driving changes to the CV_SS_ of the population size distribution, we next sought to better quantify this asymmetry. To do this, we use the Jensen-Shannon divergence *D*_*JS*_(*D, M*), where the arguments D and M indicate the probability distributions for daughter and mother sizes respectively. This measure is a symmetrized version of the Kullback-Leibler divergence that allows us to measure the level of overlap or similarity between two probability distributions (Lin, 1991). The closer *D*_*JS*_ is to 0, the more they overlap, and the closer it is to 1, the more separated they are. We then proceeded to look at how CV_SS_ correlated with such Jensen-Shannon divergence in multiple conditions, including random parameter perturbations of equivalent magnitude (**Fig. 4D**). As expected, the maximum CV_SS_ variations are found in the direction of the gradient, but, remarkably, scatter plots of the CV_SS_ as a function of the asymmetry between subpopulations show them to be strongly correlated in general. The trend is also consistent with the *experimental* values for the *cln3Δ* and *cln3Δwhi5Δ* mutant cells we examined earlier. This suggests that the differences in size between the mother and daughter populations is the dominating mode driving CV_SS_ variation. There are only a few outliers to this general trend, including the G1 timer that leads to lower asymmetry with higher CV_SS_. However, removing all G1 size control so that the transition to S phase is governed by a timer is an extreme perturbation that is not a modulation of the model parameters. Overall, these results suggest changing the level of size asymmetry between daughters and mothers is the most efficient way to modulate the CV_SS_.

### Parameter sensitivity analysis identifies subpopulations resulting from division asymmetry as a major determinant of cell size variability at *Start*

After analyzing the determinants of the CV_SS_ of the population size distribution, we conducted a similar analysis to identify which parameters influence cell size variability at *Start*. This is particularly relevant because size control at *Start* could be thought to ensure that cells reach a critical size before initiating DNA replication, rather than merely reducing CV across the population. We therefore examined the parameter gradient having the largest effect on the CV of the daughter size distribution at *Start, CV*_*Start*_. The parameter gradient is dominated by a single leading component, the division rate of mother cells, and 7 other significant components that alter the cell cycle in subtle ways (**Fig. 4A**). Perturbing the model along the parameter gradient directly affects the size asymmetry at *Start* and has a much larger effect on CV_*Start*_ than random parameter perturbations (**Fig. 4E-F**). Perturbations along the gradient increase the size asymmetry between daughters born from first-generation cells and those born from second-generation or later cells (**Fig. 4E**). As previously noted, due to the exponential accumulation of size, daughters born from larger mother cells are themselves larger than those born from first-generation cells. For WT cells, this difference in size is present but much smaller. However, perturbing the model along the gradient direction accentuates the effect and greatly changes CV_*Start*_. While significant, perturbations along the gradient have smaller effects on the CV than a pure deterministic sizer, which forces all cells below the critical size to pass through *Start* at the same threshold to sharply narrow CV_*Start*_. After performing the gradient analysis, we sought to determine if there was a general connection between the subpopulation structure and CV_*Start*_. To do this, we again used the Jensen-Shannon divergence, but this time to measure the differences between the size distributions of the daughters born from 1st generation cells and the daughters born from 2nd generation or later cells, *D*_*JS*_(*D*_1_, *D*_2+_) (**Fig. 4G**). This revealed a clear correlation between CV_*Start*_ and the daughter cell population structure. Outliers of the trend correspond to drastic parameter changes, such as specifying a deterministic sizer or timer mechanism.

It is important to note that our parameter sensitivity analysis assumes all 21 parameters of the model are independent from each other. In live cells however, there is coordination between cell cycle events meaning that there probably exist correlations between some of our 21 parameters that might be revealed by a larger scale mutational analysis. Thus, when we define a pure sizer or timer in pre-*Start* G1 (**Fig. 2C,D**) the parameters regulating the rest of the cell cycle remain unchanged. This could explain the discrepancy between the observed asymmetry of the *cln3Δwhi5Δ* mutant and the pre-*Start* G1 timer models as shown in **Fig. 4D**. Both models display a similar CV, but the asymmetry between the daughter and the mother sub-distributions is decreased rather than increased, as our analysis would predict.

### Mother-daughter asymmetry is highly correlated with population CV in *S. cerevisae* cells growing on different carbon sources

Our computational analysis suggests that mother-daughter asymmetry is a key determinant of the CV of cell size. However, testing this hypothesis experimentally was not straightforward because the mutations we have previously studied primarily affect the *Start* transition and do not directly alter mother-daughter asymmetry. To address this, we need to find a way to increase mother-daughter asymmetry while minimally impacting other cell-cycle aspects. We thus examined WT cells growing under different conditions. This is a ‘natural’ perturbation, minimally perturbating cell homeostasis, where we further expected that changes of growth rates would give rise to variations in mother-daughter size-asymmetry. This expectation is based on multiple findings. First, the duration of the budded phase, when cell growth is directed into the daughter, remains relatively constant across different growth rates (Hartwell & Unger, 1977). Because the daughter-to-mother size ratio at division is determined by the length of the budded phase relative to the time required to double a cell’s biomass, slower growth should increase mother-daughter asymmetry. Second, the growth rate is one of the dominant parameters of the CV_ss_ gradient direction as identified by our parameter sensitivity analysis (**Fig. 4A**). Thus, by analyzing yeast cells in different growth conditions, we could experimentally test our hypothesis that mother-daughter asymmetry is a primary determinant of the CV of cell size.

To generate different cell size distributions, we examined *S. cerevisiae* cells growing in synthetic complete media with 12 different carbon sources using a Coulter counter to measure cell size (see **Materials and Methods**; **Fig. 5A**). These distributions exhibited a range of cell size CVs from 0.4 to 0.62 (CV_WT_ = 0.41), which largely, but not completely, correlated with the population growth rate (R^2^=0.6; **Fig. 5B**). This is consistent with the fact that growth rate indeed is a major determinant of CV_ss_.

**Figure 5:**
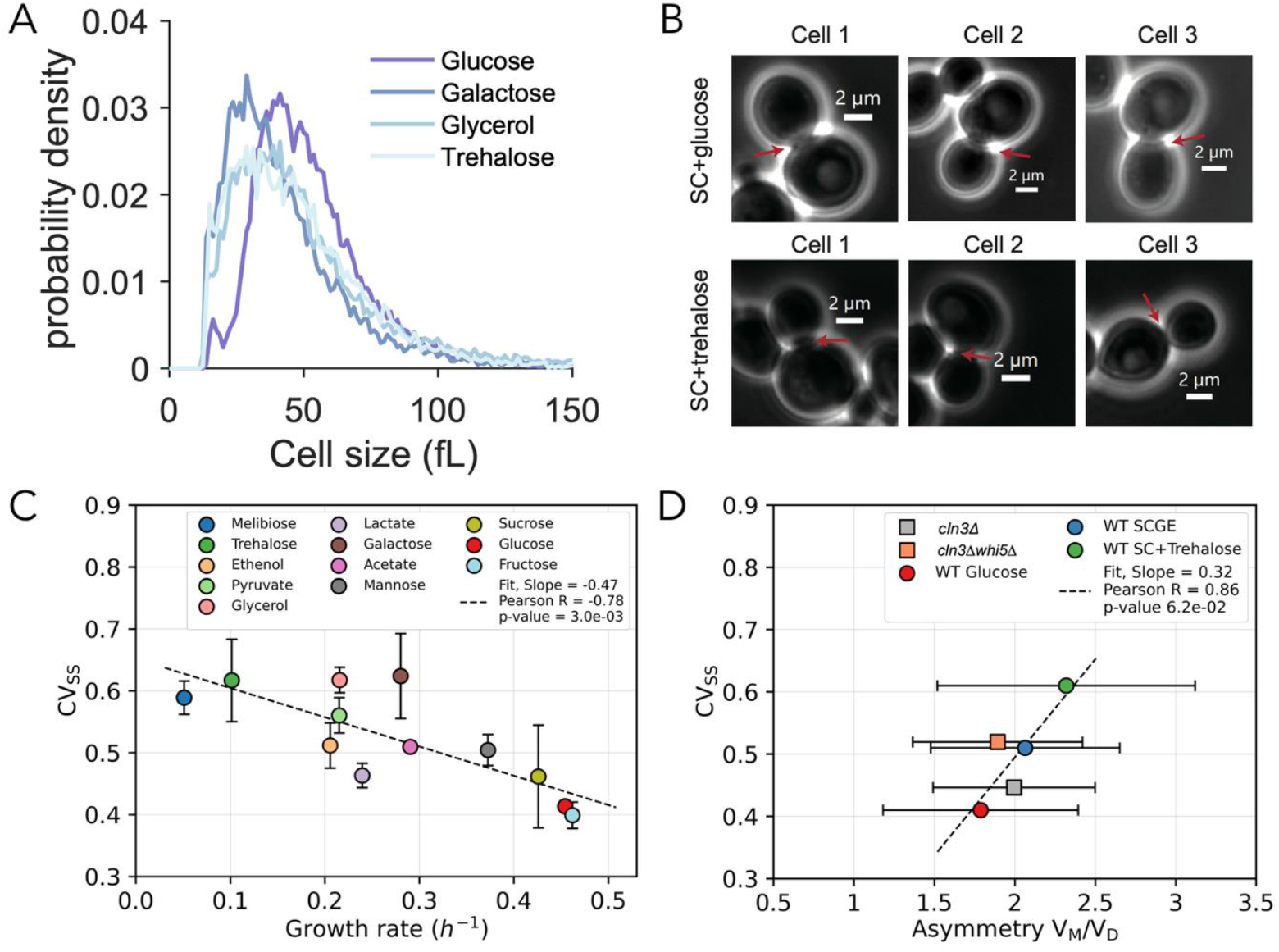
Mother-daughter asymmetry is highly correlated with population CV for *S. cerevisae* cells growing in different nutrient conditions. **(A)** Steady-state cell size distributions for *S. cerevisae* cells growing in the indicated nutrient condition. **(B)** Composite phase and fluorescence images of cells dividing into smaller daughter and larger mother cells in the indicated nutrient conditions. **(C)** CV of the steady state cell size distribution plotted as a function of the exponential growth rate for cells growing in that condition, computed from Coulter Counter distributions. The growth rate was altered by growing cells in synthetic complete media with different carbon sources. Error bars indicate standard deviation of measurements for 2 or 3 replicates. **(D)** CV at steady state plotted against the median of the volume asymmetry between pairs of mother cells (V_M_) and daughter cells (V_D_) at birth, using microscopy data allowing to monitor cell pedigree (one experiment per condition). Horizontal error bars indicate the standard deviation of the asymmetry ratio at division for cells growing in a given nutrient condition of a given genotype.

Next, we sought to measure the asymmetry at division in several different growth conditions exhibiting a range of CVs. To do this, we examined *MYO1-mKate* cells that express a fluorescent Myo1-mKate fusion protein from its endogenous locus. This myosin protein disappears from the bud neck following cytokinesis to allow an accurate determination of the time of division (Bi et al., 1998), and the corresponding daughter-to-mother size ratio (**Fig. 5C**). Consistent with our hypothesis, we find a strong correlation (R^2^=0.86) between the daughter-to-mother size ratio and the population CV (**Fig. 5D**). This confirms the model prediction that daughter-to-mother size ratio is a major mechanism that drives CV_ss_ variation.

## Discussion

Our study analyzes the determinants of cell size variability in *S. cerevisiae* to identify mother-daughter asymmetry as a primary driver of the coefficient of variation (CV) of cell size within a population. While previous research has largely focused on size control mechanisms at *Start*, we find that even the strongest possible perturbations to the G1/S transition have a limited impact on CV, typically in the range of 10-25%. Instead, the degree of size asymmetry at division plays a dominant role in shaping population-wide size variability. This conclusion is supported by both computational modeling and experimental measurements, which reveal a strong correlation between mother-daughter size asymmetry and CV across different genetic and environmental conditions. These findings challenge the prevailing view that G1 size control alone is sufficient to regulate population size variability and instead suggest that asymmetric division is integral to cell size homeostasis. Indeed, division asymmetry poses a significant barrier to maintaining a narrow cell size distribution.

Our computational parameter sensitivity analysis further refines this understanding by identifying two distinct types of asymmetry that influence CV: (1) the size difference between mother and daughter cells, and (2) the differential size of daughters born from first-generation and later-generation mothers. These asymmetries emerge from the growth dynamics of larger mother cells producing larger daughters, due to exponential biomass accumulation, and mother cells growing larger with each cell division cycle. Experimentally, we validate these predictions by growing yeast in different nutrient conditions and observing that variations in growth rate correlate with shifts in mother-daughter size asymmetry and, consequently, in CV. These results underscore that size asymmetry is not merely a byproduct of the asymmetric cell division cycle of budding yeast but is the key feature controlling cell size heterogeneity.

More broadly, our findings suggest that population-level size control in budding yeast arises from the interplay between asymmetric division and size control at *Start*, rather than size control alone. While deterministic sizer mechanisms at *Start* can narrow the size distribution, they do not significantly alter the underlying size asymmetry between subpopulations and therefore have limited effect on the population CV. However, sizer mechanisms at *Start* do have a *crucial* effect on the cell size at the time of DNA replication and to ensure a certain minimal size before attempting cell division. Ultimately, while we do not currently know the evolutionary pressures that shape the asymmetric division cycle of budding yeast and other species, we can certainly say that it poses a significant problem for maintaining a narrow distribution of cell sizes.

Finally, that mother-daugher asymmetry is strongly correlated with CV is in line with the existence of a small number of ‘soft-modes’ (or in machine learning/data science term, low dimensional reduction) observed in multiple biological systems (Russo et al., 2025). It has been proposed that such soft-modes naturally evolve to allow for homeostatic control. It is much ‘easier’ biologically speaking to control a few canalized modes than to independently buffer the potentially infinite number of possible perturbations. Our simulations and experiments are consistent with the existence of a soft-mode in cell size distributions for two reasons. First, random perturbations (both in models and experimentally) appear to move the system along a dominating direction of increased mother-daughter asymmetry strongly correlated with CV_SS_. Second, most random mutations nevertheless have a limited effect on CV_SS_ (Chen et al., 2020). This can further be seen and explained with the Chandler-Brown model. Namely, most random mutations barely change CV_SS_ (10% or less for most perturbations, compared to up to 40% for directed perturbations of equivalent magnitude), precisely because a random parameter change in a high-dimension parameter space has very little chance to be significantly aligned with the dominating direction of variation (the soft-mode). Taken together, our analysis shows how combining modelling and experiments to identify gradients of variations of phenotypes can uncover controllable and interpretable soft-modes, such as mother-daughter division asymmetry.

## Materials and Methods

### Mathematical Model

The Chandler-Brown (Chandler-Brown et al., 2017) model simulates a population of cell volumes growing exponentially with time. We note here that we use volume interchangeably with the total protein content, which was found to accumulate exponentially. Geometric volume exhibited slightly different dynamics, possibly due to systematic error in measurement of the complex geometric dynamics that characterize the budding yeast cell cycle and possibly due to modest changes in protein density. After birth, there are separate cell cycles for the smaller daughter and larger mother cells (**Fig. 2A**). The daughter cells begin their cell cycle at birth in pre-*Start* G1. Then, at each time step, the cell has a chance of progressing through the *Start* checkpoint with a rate *k*_*Start*_ which is a function of the current cell size. In the original study, the authors performed logistic regressions on various predictors to identify the best statistical predictor for the *Start* transition, which led them to identify cell size as the single best predictor. For this reason, the authors chose to model this size-dependent rate of going through the *Start* checkpoint by a linear function which has two parameters, the slope *α* and the offset *V*_0_ (**Fig. 2B**). That the rate of progression through *Start* increases with cell size V is the model’s implementation of G1 cell size control in daughter cells. This results in the time spent in G1 being inversely correlated with cell size at birth, as previously observed (Di Talia et al., 2007). After *Start*, daughter cells enter a short post-*Start* G1 timer period where cells spend roughly the same amount of time in this phase regardless of their size at the *Start* transition. This behavior is implemented through the very weak size dependence of the stochastic rate of progression to the next phase of the cycle (see **Fig. 2D** for a different but similar example). Next, cells enter the final S/G2/M phase of the cell cycle in which all cell growth is directed into the bud that will become the next daughter cell at division. At each time step during S/G2/M, cells have a chance of dividing proportional to the stochastic rate *k*_*Div*_, which is a linear function of the current cell size and the cell size at *Start*. Importantly, just like for the post-Start G1 transition rate, *k*_*Div*_ is only weakly dependent on cell size and S/G2/M is close to a timer as well. After division, the bud compartment and the main compartment are separated, and the bud begins a new cell cycle as a daughter cell, while the mother begins its separate cell cycle as a second-generation mother cell. As mother cells are born bigger and display little to no amount of cell size control, they are modelled as beginning their cell cycle already at the *Start* transition. Mother cells spend a short amount of time in post-*Start* G1 before entering an S/G2/M characterized by a stochastic rate *k*_*Div*_. Daughter and mother cell cycles are molecularly different, as there are several daughter specific proteins including Ace2 and Ash1 that impact the cell cycle (Colman-Lerner et al., 2001; Di Talia et al., 2009). This is implemented in the model by daughter cells and mother cells having independent and different parameter values for their cell cycle progression rates but having the same functional forms.

The model simulates an exponentially growing population of thousands of cells independently going through their cell cycles, but does not include cell death. For numerical purposes, we allow a maximal number of cells in the population such that when this number is reached, cells are randomly dropped out of the population. Over time, this allows us to reach a steady state in the growth of the population and allows us to look at the cell size distribution at various points of the cell cycle.

To generate the model predictions of Figures 2 and 4, we simulated the cell cycles of a maximum of 10^4^ cells for 2000 time-steps corresponding to an equivalent number of minutes for cell growth and cell cycle progression. In all simulations, we ensured the population reached steady state before performing the analysis. We refer the reader to the original publication of the model for more details on the implementation (Chandler-Brown et al., 2017).

### Parameter sensitivity analysis

To compute derivatives of any function f(*θ*) with respect to the logarithm of the parameters, we simply perform a change of variable to find that 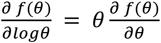 Using this equality, we can simply use numerical differentiation to approximate the gradient along the parameter direction *θ* with the following formula 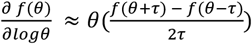, where τ is the differentiation step. We can take *τ* to be proportional to the value of *θ* such that *τ* = *hθ*, where h is a small relative perturbation (we choose h = 0.05). Thus, we get the final equation for the component of the gradient along parameter direction *θ* with respect to the logarithm of the parameter:

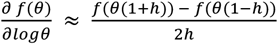

Each gradient component computation requires running the model twice. First, we run the model with the corresponding parameter component altered to *θ*(1 + *h*), and then a second time with *θ*(1 − *h*). For each of these runs, we let the population reach steady-state and compute the size distributions from which we extract mean, standard deviation and CV. These measures are then used in the equation above to approximate the gradient component. We then repeat this computation for all 21 independent parameter directions.

### Experimental strains and growth conditions

The XG04 strain was used to measure the coefficient of variation of cell size distributions in different growth conditions. The *MYO1-mKate* strain (MS358) was used to determine daughter and mother cell size upon division by microscopy. Yeast cells were cultured in synthetic complete media (SC) with 2% different carbon sources. For the microscopy experiments, SC + 2% glucose, SC + 2% glycerol + 1% ethanol, SC + 2% trehalose were used. For the coefficient of variation measurements, SC media with 2% different carbon sources (fructose, glucose, sucrose, mannose, acetate, galactose, lactate, ethanol, pyruvate, glycerol, trehalose, and melibiose) were used.

### Cell growth rate measurements

To measure the growth rate of cells in a specific carbon source, cells were first cultured for more than 10 doubling times to ensure they were in the steady state of the exponential phase. Cells were then diluted into a pre-heated flask with 50 ml of fresh media (OD600=0.01). Cells were incubated at 30°C and OD600 was measured every 1-2 hours. To determine the specific growth rate, the slope of log (OD600) versus time was calculated using a linear fit. There were at least two biological replicates for each condition.

### Coefficient of variation measurements

Cells were first cultured for more than 10 doubling times to ensure they were in the steady state of the exponential phase. Cells growing at an OD600 of 0.1–0.2 were then sonicated and diluted 100 times in 10 mL of Isoton II diluent (Beckman Coulter #8546719) prior to measurement. The coefficient of variation was determined using a Beckman Coulter Z2 counter. Cell volumes below 10 femtoliters (fL) or above 300 fL were excluded from the analysis. There were at least two biological replicates for each condition.

### Mother-to-daughter cell size ratio measurements

Cells expressing a Myo1-mKate myosin fusion protein were used to determine the cell size of mother and daughter cells at division. Yeast cells were grown in SC media supplemented with the indicated carbon sources to exponential phase (OD = 0.1–0.2). 80 µl cell culture and 200 µl fresh medium were loaded into the corresponding wells of a microfluidic plate (CellASIC #Y04C-02-5PK). The microfluidic plate was then connected to the CellASIC ONIX2 Microfluidic System to control the media flow rates. Media was initially perfused into the chamber for 2 minutes at 8 psi, followed by cell loading for 10 s at 2 psi. Continuous perfusion of fresh medium at 2 psi was maintained until imaging was completed. A Tempcontrol 37-2 digital system ensured cell growth at 30°C. Cells were imaged on a wide-field epifluorescence Zeiss AXIO Observer.Z1 microscope (63X/1.4NA oil immersion objective and a Colibri LED module). mKate was imaged in the red channel (555 nm LED module; 25% intensity; 1 second exposure) and phase contrast images were taken to give cell boundaries. Cells in glucose and glycerol/ethanol were imaged for 8 hours at 5 min intervals, while cells in trehalose were imaged for 15 hours at 10 min intervals. Cell boundaries were automatically segmented in Cell-ACDC (Padovani et al., 2022) using the software’s default settings, followed by manual correction of segmentation errors. For S/G2/M phase cells, the disappearance of the MYO1-mKate signal at the bud neck marked the time of the cell division. At division, mother and daughter cell sizes were then obtained using Cell-ACDC’s volume algorithm.

#### Yeast strains

MS358

W303 *WHI5pr-WHI5-mCitrine::ura; whi5Δ::cgTRP; MYO1-3xmKate2::kanMX6; ADE2 MAT****a***

Source Swaffer et al., 2023

XG04

W303 *PUS1-EGFP:KanM*6*; *pdr5Δ:leu2; RPS9A-Halotag:HygB; ADE2 MAT****a***

Source This study

## Bibliography

Amir, A. (2014). Cell Size Regulation in Bacteria. Physical Review Letters, 112(20), 208102. 10.1103/PhysRevLett.112.208102

Barber, F., Amir, A., & Murray, A. W. (2020). Cell-size regulation in budding yeast does not depend on linear accumulation of Whi5. Proceedings of the National Academy of Sciences, 117(25), 14243–14250. 10.1073/pnas.2001255117

Berenson, D. F., Zatulovskiy, E., Xie, S., & Skotheim, J. M. (2019). Constitutive expression of a fluorescent protein reports the size of live human cells. Molecular Biology of the Cell, 30(24), 2985–2995. 10.1091/mbc.E19-03-0171

Bi, E., Maddox, P., Lew, D. J., Salmon, E. D., McMillan, J. N., Yeh, E., & Pringle, J. R. (1998). Involvement of an Actomyosin Contractile Ring in Saccharomyces cerevisiae Cytokinesis. Journal of Cell Biology, 142(5), 1301–1312. 10.1083/jcb.142.5.1301

Bruin, R. A. M. de, McDonald, W. H., Kalashnikova, T. I., Yates, J., & Wittenberg, C. (2004). Cln3 Activates G1-Specific Transcription via Phosphorylation of the SBF Bound Repressor Whi5. Cell, 117(7), 887–898. 10.1016/j.cell.2004.05.025

Chandler-Brown, D., Schmoller, K. M., Winetraub, Y., & Skotheim, J. M. (2017). The Adder Phenomenon Emerges from Independent Control of Pre- and Post-Start Phases of the Budding Yeast Cell Cycle. Current Biology, 27(18), Article 18. 10.1016/j.cub.2017.08.015

Chen, Y., Zhao, G., Zahumensky, J., Honey, S., & Futcher, B. (2020). Differential Scaling of Gene Expression with Cell Size May Explain Size Control in Budding Yeast. Molecular Cell, 78(2), Article 2. 10.1016/j.molcel.2020.03.012

Colman-Lerner, A., Chin, T. E., & Brent, R. (2001). Yeast Cbk1 and Mob2 Activate Daughter-Specific Genetic Programs to Induce Asymmetric Cell Fates. Cell, 107(6), 739–750. 10.1016/S0092-8674(01)00596-7

Costanzo, M., Nishikawa, J. L., Tang, X., Millman, J. S., Schub, O., Breitkreuz, K., Dewar, D., Rupes, I., Andrews, B., & Tyers, M. (2004). CDK Activity Antagonizes Whi5, an Inhibitor of G1/S Transcription in Yeast. Cell, 117(7), 899–913. 10.1016/j.cell.2004.05.024

Cross, F. R. (1988). DAF1, a Mutant Gene Affecting Size Control, Pheromone Arrest, and Cell Cycle Kinetics of Saccharomyces cerevisiae. Molecular and Cellular Biology, 8(11), 4675–4684. 10.1128/mcb.8.11.4675-4684.1988

Crozier, L., Foy, R., Adib, R., Kar, A., Holt, J. A., Pareri, A. U., Valverde, J. M., Rivera, R., Weston, W. A., Wilson, R., Regnault, C., Whitfield, P., Badonyi, M., Bennett, L. G., Vernon, E. G., Gamble, A., Marsh, J. A., Staples, C. J., Saurin, A. T., … Ly, T. (2023). CDK4/6 inhibitor-mediated cell overgrowth triggers osmotic and replication stress to promote senescence. Molecular Cell, 83(22), 4062-4077.e5. 10.1016/j.molcel.2023.10.016

Di Talia, S., Skotheim, J. M., Bean, J. M., Siggia, E. D., & Cross, F. R. (2007). The effects of molecular noise and size control on variability in the budding yeast cell cycle. Nature, 448(7156), 947–951. 10.1038/nature06072

Di Talia, S., Wang, H., Skotheim, J. M., Rosebrock, A. P., Futcher, B., & Cross, F. R. (2009). Daughter-Specific Transcription Factors Regulate Cell Size Control in Budding Yeast. PLoS Biology, 7(10), e1000221. 10.1371/journal.pbio.1000221

Eun, Y.-J., Ho, P.-Y., Kim, M., LaRussa, S., Robert, L., Renner, L. D., Schmid, A., Garner, E., & Amir, A. (2018). Archaeal cells share common size control with bacteria despite noisier growth and division. Nature Microbiology, 3(2), 148–154. 10.1038/s41564-017-0082-6

Fantes, P. A. (1977). Control of cell size and cycle time in Schizosaccharomyces pombe. Journal of Cell Science, 24(1), 51–67. 10.1242/jcs.24.1.51

Foy, R., Crozier, L., Pareri, A. U., Valverde, J. M., Park, B. H., Ly, T., & Saurin, A. T. (2023). Oncogenic signals prime cancer cells for toxic cell overgrowth during a G1 cell cycle arrest. Molecular Cell, 83(22), 4047-4061.e6. 10.1016/j.molcel.2023.10.020

Ginzberg, M. B., Kafri, R., & Kirschner, M. (2015). Cell biology. On being the right (cell) size. Science (New York, N.Y.), 348(6236), 1245075. 10.1126/science.1245075

Gutenkunst, R. N., Waterfall, J. J., Casey, F. P., Brown, K. S., Myers, C. R., & Sethna, J. P. (2007). Universally Sloppy Parameter Sensitivities in Systems Biology Models. PLOS Computational Biology, 3(10), e189. 10.1371/journal.pcbi.0030189

Hartwell, L. H., & Unger, M. W. (1977). Unequal division in Saccharomyces cerevisiae and its implications for the control of cell division. The Journal of Cell Biology, 75(2 Pt 1), 422–435. 10.1083/jcb.75.2.422

Johnston, G. C., Pringle, J. R., & Hartwell, L. H. (1977). Coordination of growth with cell division in the yeast Saccharomyces cerevisiae. Experimental Cell Research, 105(1), 79–98. 10.1016/0014-4827(77)90154-9

Jun, S., Si, F., Pugatch, R., & Scott, M. (2018). Fundamental principles in bacterial physiology— history, recent progress, and the future with focus on cell size control: A review. Reports on Progress in Physics, 81(5), Article 5. 10.1088/1361-6633/aaa628

Khurana, A., Chadha, Y., & Schmoller, K. M. (2023). Too big not to fail: Different paths lead to senescence of enlarged cells. Molecular Cell, 83(22), 3946–3947. 10.1016/j.molcel.2023.10.024

Lanz, M. C., Zatulovskiy, E., Swaffer, M. P., Zhang, L., Ilerten, I., Zhang, S., You, D. S., Marinov, G., McAlpine, P., Elias, J. E., & Skotheim, J. M. (2022). Increasing cell size remodels the proteome and promotes senescence. Molecular Cell, 82(17), 3255-3269.e8. 10.1016/j.molcel.2022.07.017

Lanz, M. C., Zhang, S., Swaffer, M. P., Ziv, I., Götz, L. H., Kim, J., McCarthy, F., Jarosz, D. F., Elias, J. E., & Skotheim, J. M. (2024). Genome dilution by cell growth drives starvationlike proteome remodeling in mammalian and yeast cells. Nature Structural & Molecular Biology, 31(12), 1859–1871. 10.1038/s41594-024-01353-z

Lin, J. (1991). Divergence measures based on the Shannon entropy. IEEE Transactions on Information Theory, 37(1), 145–151. 10.1109/18.61115

Lloyd, A. C. (2013). The Regulation of Cell Size. Cell, 154(6), 1194–1205. 10.1016/j.cell.2013.08.053

Manohar, S., Estrada, M. E., Uliana, F., Vuina, K., Alvarez, P. M., de Bruin, R. A. M., & Neurohr, G. E. (2023). Genome homeostasis defects drive enlarged cells into senescence. Molecular Cell, 83(22), 4032-4046.e6. 10.1016/j.molcel.2023.10.018

Miettinen, T. P., & Björklund, M. (2016). Cellular Allometry of Mitochondrial Functionality Establishes the Optimal Cell Size. Developmental Cell, 39(3), Article 3. 10.1016/j.devcel.2016.09.004

Nash, R., Tokiwa, G., Anand, S., Erickson, K., & Futcher, A. B. (1988). The WHI1+ gene of Saccharomyces cerevisiae tethers cell division to cell size and is a cyclin homolog. The EMBO Journal, 7(13), 4335–4346. 10.1002/j.1460-2075.1988.tb03332.x

Padovani, F., Mairhörmann, B., Falter-Braun, P., Lengefeld, J., & Schmoller, K. M. (2022). Segmentation, tracking and cell cycle analysis of live-cell imaging data with Cell-ACDC. BMC Biology, 20(1), 174. 10.1186/s12915-022-01372-6

Qu, Y., Jiang, J., Liu, X., Wei, P., Yang, X., & Tang, C. (2019). Cell Cycle Inhibitor Whi5 Records Environmental Information to Coordinate Growth and Division in Yeast. Cell Reports, 29(4), 987-994.e5. 10.1016/j.celrep.2019.09.030

Russo, C. J., Husain, K., & Murugan, A. (2025). Soft Modes as a Predictive Framework for Low-Dimensional Biological Systems Across Scales. Annual Review of Biophysics, 54(Volume 54, 2025), 401–426. 10.1146/annurev-biophys-081624-030543

Schmoller, K. M., Lanz, M. C., Kim, J., Koivomagi, M., Qu, Y., Tang, C., Kukhtevich, I. V., Schneider, R., Rudolf, F., Moreno, D. F., Aldea, M., Lucena, R., & Skotheim, J. M. (2022). Whi5 is diluted and protein synthesis does not dramatically increase in pre-Start G1. Molecular Biology of the Cell, 33(5), lt1. 10.1091/mbc.E21-01-0029

Schmoller, K. M., Turner, J. J., Kõivomägi, M., & Skotheim, J. M. (2015). Dilution of the cell cycle inhibitor Whi5 controls budding-yeast cell size. Nature, 526(7572), 268–272. 10.1038/nature14908

Sveiczer, A., Novak, B., & Mitchison, J. M. (1996). The size control of fission yeast revisited. Journal of Cell Science, 109(12), Article 12. 10.1242/jcs.109.12.2947

Tzur, A., Kafri, R., LeBleu, V. S., Lahav, G., & Kirschner, M. W. (2009). Cell Growth and Size Homeostasis in Proliferating Animal Cells. Science, 325(5937), 167–171. 10.1126/science.1174294

Wang, H., Carey, L. B., Cai, Y., Wijnen, H., & Futcher, B. (2009). Recruitment of Cln3 Cyclin to Promoters Controls Cell Cycle Entry via Histone Deacetylase and Other Targets. PLOS Biology, 7(9), e1000189. 10.1371/journal.pbio.1000189

Westfall, C. S., & Levin, P. A. (2017). Bacterial Cell Size: Multifactorial and Multifaceted. Annual Review of Microbiology, 71, 499–517. 10.1146/annurev-micro-090816-093803

Willis, L., & Huang, K. C. (2017). Sizing up the bacterial cell cycle. Nature Reviews Microbiology, 15(10), 606–620. 10.1038/nrmicro.2017.79

Xie, S., & Skotheim, J. M. (2020). A G1 Sizer Coordinates Growth and Division in the Mouse Epidermis. Current Biology: CB, 30(5), 916-924.e2. 10.1016/j.cub.2019.12.062

Xie, S., Swaffer, M., & Skotheim, J. M. (2022). Eukaryotic Cell Size Control and Its Relation to Biosynthesis and Senescence. Annual Review of Cell and Developmental Biology, 38, 291–319. 10.1146/annurev-cellbio-120219-040142

Xie, S., Zhang, S., Medeiros, G. de, Liberali, P., & Skotheim, J. M. (2024). The G1/S transition in mammalian stem cells in vivo is autonomously regulated by cell size (p. 2024.04.09.588781). bioRxiv. 10.1101/2024.04.09.588781

Zatulovskiy, E., Lanz, M. C., Zhang, S., McCarthy, F., Elias, J. E., & Skotheim, J. M. (2022). Delineation of proteome changes driven by cell size and growth rate. Frontiers in Cell and Developmental Biology, 10. 10.3389/fcell.2022.980721

